# Estimating recombination using only the allele frequency spectrum

**DOI:** 10.1101/2025.02.01.635998

**Authors:** Matthew W. Hahn, Sarthak R. Mishra

## Abstract

Standard methods for estimating the population recombination parameter, *ρ*, are dependent on sampling individual genotypes and calculating various types of disequilibria. However, recent machine learning (ML) approaches to estimating recombination have used pooled sequencing data, which does not sample individual genotypes and cannot be used to calculate disequilibria beyond the length of a single sequence read. Motivated by these results, this study examines the “black box” of such ML methods to understand what signals are being used to infer recombination rates. We find that it is indeed possible to estimate recombination solely using the allele frequency spectrum, and we provide a genealogical interpretation of these results. We further show that even a simplified representation of the allele frequency spectrum can be used to estimate recombination. We demonstrate the accuracy of such inferences using both simulations and data from humans. These results offer a new way to understand the effects of recombination on patterns of sequence data, as well as providing an example of how the internal workings of ML methods can give insight into biological processes.

**Article Summary:** Machine learning methods are becoming more common, offering powerful approaches to study the natural world. We investigated a popular machine learning method to see how it worked, discovering that it was exploiting data (the allele frequency spectrum) to estimate genetic recombination rates that had not been considered before. Our study demonstrates that this approach is indeed quite powerful, opening up new avenues of research. The work also demonstrates that looking inside machine learning models can sometimes teach us novel things about nature.

## Introduction

Recombination is a fundamental biological process that plays an important role in evolution (Johnston 2024). While crosses between individuals and the genotyping of a large number of offspring are often used to infer the meiotic recombination rate, *c*, the population recombination parameter, *ρ* (=4*N*_e_*c*), can be inferred from a small sample of unrelated individuals. The magnitude of this parameter reflects the history of recombination in the sample across many thousands of generations, but is often strongly correlated with the underlying meiotic recombination rate (e.g. McVean et al. 2004; Stevison et al. 2016).

There are multiple common ways to estimate *ρ* (reviewed in Hahn 2018, chapter 4; Peñalba and Wolf 2020). Probably the most widely used set of methods are based on gametic linkage disequilibrium (LD), using individually phased haplotypes to estimate the association between alleles on chromosomes. Measures of gametic LD can then be used to estimate *ρ* (Sved 1971; Weir and Hill 1986; McVean 2002), or haplotypes can be used directly (e.g. Hudson 1987; Wakeley 1997; Wall 2000). If phased haplotypes are not available, another form of LD can still be calculated from diploid genotypes: genotypic LD (Weir 1979). The very popular (and accurate) class of methods that estimate *ρ* using composite likelihood (Hudson 2001; McVean et al. 2002; Chan et al. 2012; Kamm et al. 2016; Spence and Song 2019) can all use either phased haplotypes (i.e. gametic LD) or unphased genotypes (i.e. genotypic LD). Finally, a newer set of approaches based solely on whether positions are heterozygous or homozygous—without respect to the particular alleles or genotypes at a site—have been used to calculate so-called zygotic LD and consequently *ρ* (Haubold et al. 2010; Barroso et al. 2019; Setter et al. 2022). Despite the relative lack of resolution in the recombination rate using zygotic LD, such approaches are also highly accurate (Dutheil 2024).

In the past few years, machine learning (ML) methods have become a useful and accurate approach for multiple types of inference in population genetics (Schrider and Kern 2018; Korfmann et al. 2023; Huang et al. 2024). ML methods are especially useful in dealing with messy data: in the case of estimating *ρ*, this might mean incorrectly inferred haplotypes or genotypes. Indeed, multiple ML approaches for estimating *ρ* have been introduced over the past dozen years (Lin et al. 2013; Gao et al. 2016; Flagel et al. 2019; Hermann et al. 2019), including ReLERNN (Recombination Landscape Estimation using Recurrent Neural Networks), a deep learning tool that can accurately infer *ρ* from sub-optimal data (Adrion et al. 2020).

Most interestingly, ReLERNN is also able to accurately infer *ρ* from pooled sequencing data. Pooled sequencing (sometimes called “pool-seq”; Schlötterer et al. 2014) provides only allele frequencies at each genomic position, as no barcodes or labels are associated with each sampled individual in the pool. While there have been previous methods that could infer very short-range LD from pooled sequencing (Feder et al. 2012), these rely on SNPs found in the same read and are therefore limited to short distances. In contrast, ReLERNN does not take any information about sequence reads into account—the input contains only a list of SNP positions and allele frequencies within a genomic window. Although no obvious type of disequilibrium can be calculated from such data, Adrion et al. (2020) showed that ReLERNN can very accurately infer *ρ* across larger distances.

ML methods can learn from messy, high-dimensional data, but are also prone to picking up on unintended signals provided by, for instance, the order in which data are presented, seemingly innocuous data labels, or other non-meaningful aspects of the training set (Bernett et al. 2024). Putting aside the possibility of such data leakage, there are multiple signals associated with recombination that ReLERNN could be using for inferences from pooled sequencing (Adrion and colleagues do not speculate as to the source of the signal). First, the input to ReLERNN implicitly encodes the number of SNPs in a window as the number of columns in the dataset. As there is a near-universal correlation in natural populations between the number of SNPs in a region—often represented by the population mutation parameter, *θ* (=4*N*_e_*μ*)—and the population recombination parameter (Cutter and Payseur 2013), it is possible that ReLERNN could use this relationship to estimate *ρ*. However, a strong relationship between *θ* and *ρ* can only arise in non-neutral scenarios, and Adrion et al. (2020) show that their method is still accurate in neutral, equilibrium populations. Second, the input to ReLERNN contains the genomic position of each SNP in a window. While it is not obvious what sort of information the distance between variable positions might contain about recombination, it is possible that it is using this information.

Finally, and most relevant for what follows in this paper, the input to ReLERNN is comprised of the frequency of each SNP, either as the minor or derived allele frequency. A collection of allele frequencies at multiple sites can be used to construct an allele frequency spectrum, which is simply a summary of the various frequencies within a sample. Adrion et al. (2020) showed that their accuracy in estimating *ρ* increased with more accurate estimates of allele frequencies, suggesting that these data are a key input.

Here, we examine the “black box” at the heart of ML estimates of recombination from pooled sequencing data. We propose that the allele frequency spectrum can be related to the population recombination parameter, *ρ*, and we provide a genealogical explanation for this relationship. We first demonstrate this connection using simulations. We then develop a simple ML model for inferring *ρ* from pooled sequencing data, showing that it is both accurate and robust to many assumptions. We apply our model, called NoDEAR (No Disequilibrium Estimation of Accurate Recombination), to data from humans, demonstrating that it is highly correlated with estimates using composite likelihood methods. Together, our investigations shed light into novel ways that recombination can affect the allele frequency spectrum, and how ML methods can help to uncover fundamental biological relationships.

### Genealogical effects on the allele frequency spectrum

The allele frequency spectrum is a central concept in modern population genetics. For a sample of *n* haploid chromosomes, we define the allele frequency spectrum as a vector of length *n*-1 when considering derived allele frequencies and of length *n*/2 when considering minor allele frequencies (rounding down if *n* is an odd number). For the derived frequency spectrum, the entries in the vector correspond to either the count or the proportion of all polymorphisms in a dataset found on 1, 2, 3,…*n*-1 chromosomes. Elements of each vector therefore represent the fraction of all variants found at sample frequencies 1/*n*, 2/*n*, 3/*n*,…*n*-1/*n*. Such an object is perhaps not an obvious source of information about recombination. One reason for this is that the expected frequency spectrum has been derived using multiple approaches (Ewens 1979; Tajima 1989; Fu 1995; Griffiths and Tavaré 1998; Hudson 2015) and is the same across population sizes and recombination rates. However, these results are all expectations in the limit of an infinite number of loci; as we explain next, it is exactly the violation of this assumption that provides a link to recombination.

To understand how the allele frequency spectrum can tell us about recombination, consider the spectrum arising from a single non-recombining locus (Figure 1a). At a single locus there is a single tree topology, which limits the possible allele frequencies observed. Assuming an infinite sites model (i.e. the same mutation does not appear more than once), mutations can have only a limited set of frequencies: those possible in the local tree topology. Consider the toy example in Figure 1a: only polymorphisms of “size” 1 and 2 are possible, since mutations can occur only on branches with either 1 or 2 descendants (Fu 1995). Given this, only polymorphisms with a frequency of 25% or 50% are possible. In contrast, in the tree shown in Figure 1b, mutations of size 1, 2, or 3 are possible. Therefore, a different allele frequency spectrum can be produced by this tree topology.

**Figure 1.**
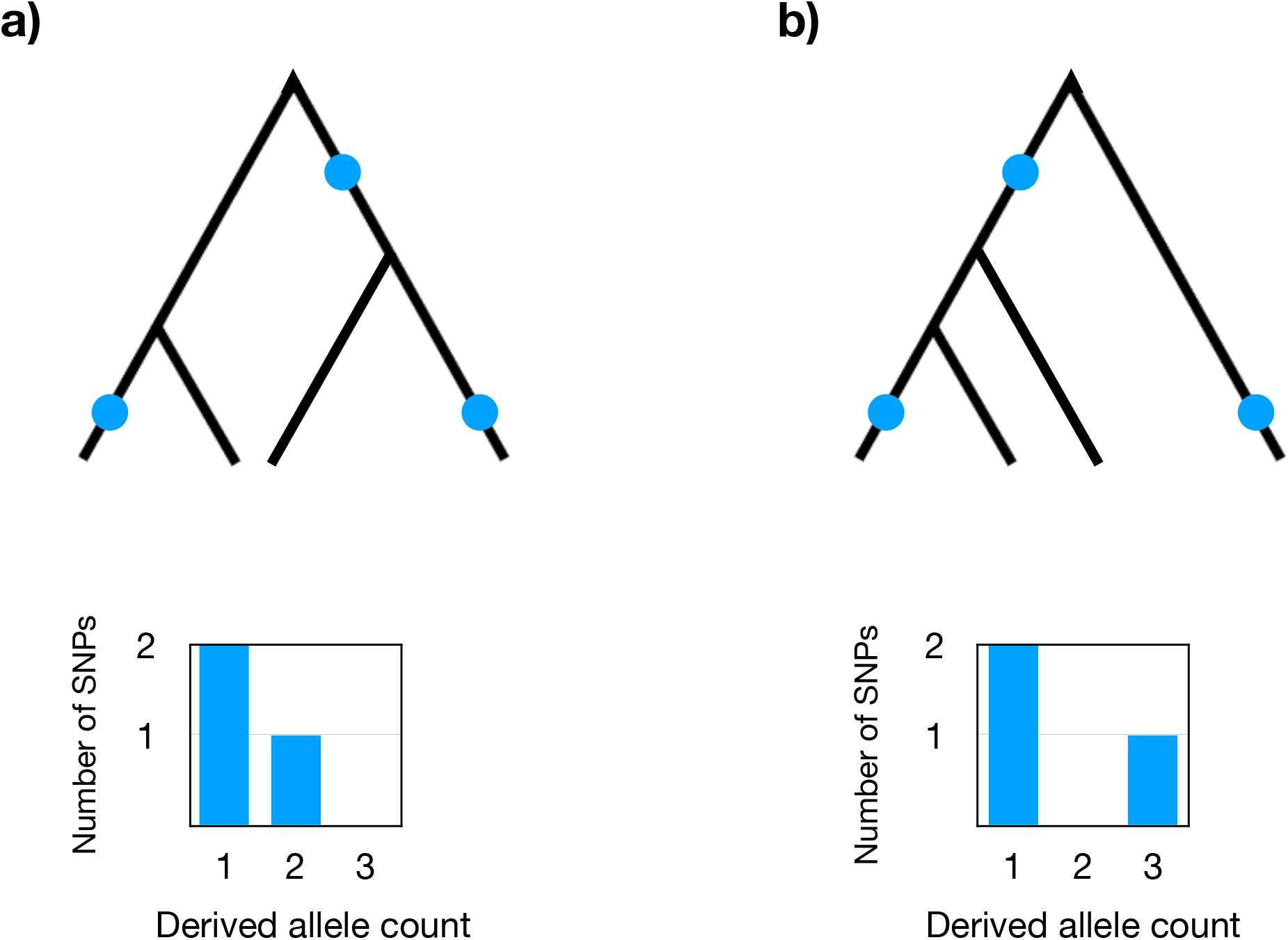
Example gene trees, mutations, and allele frequency spectra. **a)** The top shows a hypothetical gene tree with *n*=4 tips and *S*=3 mutations (blue circles). Two mutations are of size 1 (have one descendant each) and one mutation is of size 2. The bottom shows the allele frequency spectrum that would result from this tree and mutations. **b)** The top again shows a gene tree and mutations. There are two mutations of size 1 and one mutation of size 3. Importantly, under the infinite sites model, mutations of size 3 are only possible on this gene tree topology, not on the one in panel a (mutations of size 2 are also possible in this tree, but none is shown). The bottom shows the resulting allele frequency spectrum.

In general, any particular topology sampled at a non-recombining locus will permit only a subset of allele frequencies, and this subset is likely to differ among trees at different loci. Even if mutations of the same size are permitted on two trees—for instance, if they have the same hierarchical topology—differences in branch lengths can still result in different overall allele frequency spectra due to different numbers of mutations of each size (Ferretti et al. 2013). Importantly, we have proved in the Appendix (https://doi.org/10.5281/zenodo.14775487) that no single topology can possibly give the allele frequency spectrum expected in the infinite-locus limit when *n*≥4.

We propose that it is exactly the sparsity of the allele frequency spectrum from a limited number of tree topologies that provides information about recombination. Regions containing a small number of tree topologies will give sparse spectra, while regions with a large number of topologies will give smoother spectra because they are the sum across the spectra produced by each marginal topology. Every recombination event in the history of a sample will result in an additional tree, such that *R* recombination events in a genomic region leads to *R*+1 topologies in the region (Hudson 1983; Griffiths and Marjoram 1996; Wiuf and Hein 1999; McVean and Cardin 2005; Marjoram and Wall 2006). Although not all recombination events are detectable in a sample, *R* can be used to estimate *ρ* (Hudson and Kaplan 1985); this implies that the number of trees in a region is also associated with *ρ*. If the number of marginal trees in a region is related to the allele frequency spectrum in any sort of straightforward manner, we should be able to use this representation of the data to estimate *ρ*. This interpretation is also consistent with the observation by Adrion et al. (2020) that the accuracy of ReLERNN increases with higher accuracy of allele frequency estimates, as the latter will of course also make the estimate of the allele frequency spectrum more accurate.

## Results

### Simulations connect recombination to the allele frequency spectrum

To test whether a relationship between recombination and the allele frequency spectrum exists, we carried out simulations in msprime (Kelleher et al. 2016). Comprehensive simulations across parameter space were carried out as described below, here our goal was to simply demonstrate this link. We simulated *n*=20 haploid samples from the equivalent of a 50 kb region with constant population size, *N* = 70,000, and mutation rate per generation, *μ* = 1.0×10^−8^.

Figure 2 shows examples of two typical allele frequency spectra from simulations with low recombination (*ρ*=0.282) and high recombination (*ρ*=1,843). As can be seen, the spectrum from a region with less recombination in its history (Figure 2a) is bimodal, choppy, and hardly resembles the neutral expectation, while the spectrum from a region with high recombination (Figure 2b) is smoother, approaching that of the expected spectrum. For reference, we also recorded the number of unique tree topologies in the two simulations: there are 2 trees in the low recombination condition and 6,538 trees in the high recombination condition.

**Figure 2.**
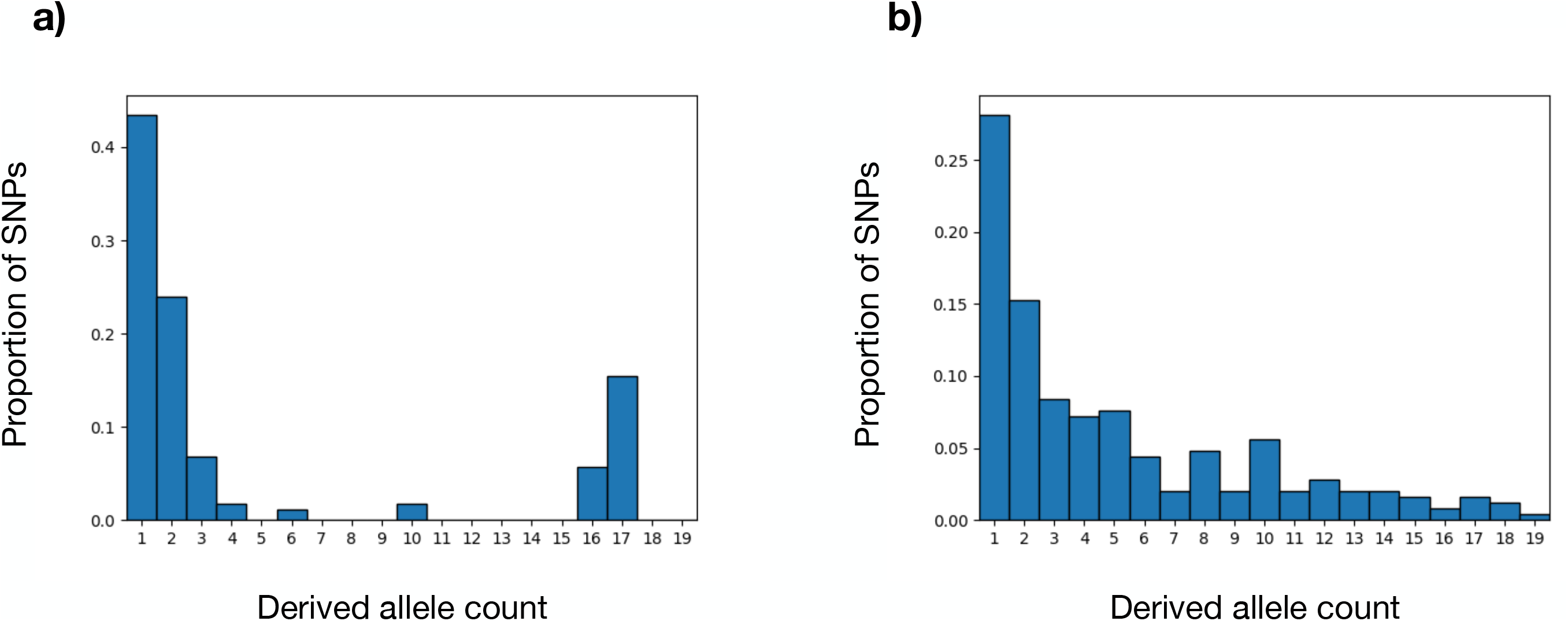
Simulated allele frequency spectra under different levels of recombination. **a)** The allele frequency spectrum produced for a sample of size *n*=20 in a simulated 50-kb region with low recombination (*ρ*=0.282). There were 2 different tree topologies found in this region. **b)** The allele frequency spectrum produced for a sample of size *n*=20 in a simulated 50-kb region with high recombination (*ρ*=1,843). There were 6,538 different tree topologies found in this region.

The qualitative descriptions of spectra as “choppy” or “smooth” are not easy to summarize across thousands of simulations and would require that we assess each allele frequency spectrum visually. Instead, we attempt to capture these patterns quantitatively by calculating the Euclidean distance (*L*^2^-norm) between the observed and expected allele frequency spectrum in each simulated window (as calculated using NumPy; Harris et al. 2020). Using this approach, genomic windows with smooth spectra will have *L*^2^-norm=0 and with highly irregular spectra will have *L*^2^-norm>>0. *L*^2^-norm can be calculated quickly for any dataset for use in quantitative comparisons. Although we use the Euclidean distance from the neutral-equilibrium expectation here, we describe how equivalent calculations can be carried out for non-equilibrium histories in the Discussion. For the examples used above, the values of *L*^2^-norm are 0.261 in low recombination (Figure 2a) and 0.049 in high recombination (Figure 2b).

Given this framework, we simulated 1000 loci with the same parameters as above, using values of *ρ* ranging across four orders of magnitude. We found a highly negative correlation between *ρ* and *L*^2^-norm calculated from the derived frequency spectrum (Spearman’s *r*_*s*_=-0.89), such that higher recombination resulted in smoother spectra and lower Euclidean distances (Supplementary Figure 1a). If we use the minor allele frequency spectrum instead, the correlation between *ρ* and *L*^2^-norm is slightly reduced (Spearman’s *r*_*s*_=-0.83; Supplementary Figure 1b); this weaker relationship is expected given that there is less information in the minor spectrum. Regardless of which we use, as predicted, the allele frequency spectrum contains information about the amount of recombination in a sample. We therefore next develop a machine learning model that can predict *ρ* from this spectrum.

### NoDEAR: a machine learning model

In light of the above relationships, we developed a simplified ML model to estimate the recombination rate using only the allele frequency spectrum. Our goal is not to compete with ReLERNN, but instead to see how little data we can use and still accurately estimate *ρ*. Here we explain how we trained our model, which we call NoDEAR (No Disequilibrium Estimation of Accurate Recombination), and following this we test its accuracy and robustness in multiple ways. The goal of NoDEAR is to highlight the power of an ML approach to recombination estimation that uses solely the allele frequency spectrum; researchers interested in estimating *ρ* from empirical datasets should use alternative software.

NoDEAR uses XGBoost (Chen and Guestrin 2016) as implemented in Python to learn relationships between the allele frequency spectrum and *ρ*. The standard input to NoDEAR is the allele frequency spectrum represented as a vector of normalized proportions, such that the sum of all entries equals 1. By normalizing the values, we remove information about the number of SNPs in a dataset. This representation also obviously contains no information about the relative genomic positions of SNPs. The output of NoDEAR is a predicted value of *ρ*.

Training our ML model was straightforward. The reference simulations were again carried out using msprime (Kelleher et al. 2016) as described in the previous section. Later, we change the population history and size of genomic windows to test the effects of each. For training, we ran simulations with *c* varying across five orders of magnitude (from 0.5×10^−12^ to 0.5×10^−7^), with 1,000 simulated datasets in each of 10 equally sized bins of increasing values of *c*, for a total of 10,000 simulated datasets. For each simulated locus, we recorded the number of recombination events, *R*, then calculated *ρ* by using the relationship *ρ* =*R/a*, where *a* is the harmonic series from 1 to *n*-1 (Hudson and Kaplan 1985). The value of *ρ* and the allele frequency vector for each locus were passed to XGBoost for training.

Training on all 10,000 simulated datasets took 45.04 seconds on one core of an Intel Xeon processor with 100 Gb of available RAM, run on the Indiana University Research Desktop. We assessed the accuracy of the NoDEAR model on the training data by carrying out five-fold cross-validation (using Scikit-learn; Pedregosa et al. 2011), with an average score of 0.917.

### Accuracy of recombination estimation from the allele frequency spectrum

After training NoDEAR on allele frequency spectra associated with a wide range of recombination histories, we asked how well this approach could predict *ρ* in new simulations not used in training. Reference simulations at 120 loci were carried out as described above, but with *c* varying across only three orders of magnitude (from 0.5×10^−11^ to 0.5×10^−8^), and then passed to NoDEAR for prediction. Runtime for prediction on the entire test dataset was 0.53 seconds. The correlation between the true value of *ρ* and the value estimated by NoDEAR was quite high, with Spearman’s *r*_*s*_=0.923 (Figure 3a; calculated using SciPy; Virtanen et al. 2020). The predicted values of *ρ* also explain a large amount of the variation in the true values of *ρ* (*R*^2^=0.813).

**Figure 3.**
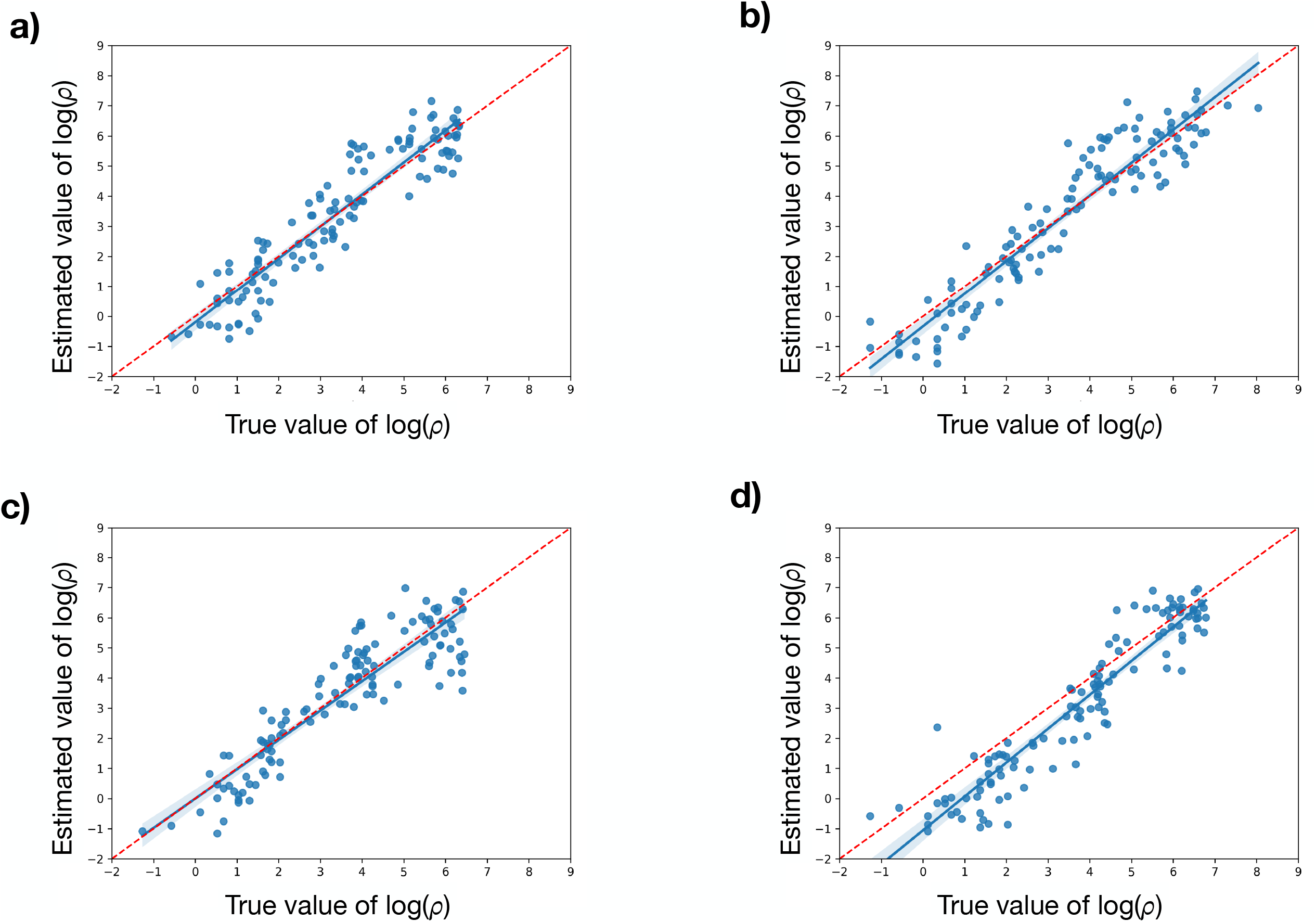
Correlation between true values of *ρ* and values estimated by NoDEAR. **a)** Correlation between values simulated under an equilibrium population history with no gene conversion (Spearman’s *r*_*s*_=0.923 and *R*^2^=0.813). Each dot represents one simulated 50-kb region. Blue lines represent best-fit regressions (as calculated in Seaborn; Waskom 2021), plus confidence intervals. Red dashed lines show the *y*=*x* line. **b)** Correlation between values simulated under non-equilibrium population histories with no gene conversion (Spearman’s *r*_*s*_=0.932; *R*^2^=0.841). **c)** Correlation between values simulated under a history with population structure and no gene conversion (Spearman’s *r*_*s*_=0.870; *R*^2^=0.778). **d)** Correlation between values simulated under an equilibrium population history with gene conversion (Spearman’s *r*_*s*_=0.928; *R*^2^=0.738)

To provide some context as to how accurate NoDEAR is compared to the state-of-the-art estimates of population recombination parameters, we applied the composite likelihood method implemented in the program, pyrho (Spence and Song 2019). We provided pyrho with the phased haplotypes from all 20 simulated individuals at each 50-kb locus, as it cannot use just the allele frequency spectrum. Pyrho outputs the recombination rate per base per generation, *c*, rather than *ρ* (=4*N*_e_*c*), by assuming that the demographic model inferred by this method is the only cause of genealogical heterogeneity (see Discussion). Regardless, the rank correlation between the inferred *c* from pyrho and the true *ρ* should be comparable to our results with NoDEAR.

Using pyrho to estimate parameters using the same test data, the correlation with the true values was extremely high, with Spearman’s *r*_*s*_=0.97 (Supplementary Figure 2; we first removed estimates of *c* below 10^−20^, as there was a cluster of outlier points with *c*≈10^−50^). Although these results are more accurate than those using NoDEAR, they also took much longer to estimate (runtime=2,549.4 seconds). In addition, increasing the sample size used by NoDEAR to *n*=100 further increases the accuracy of inference (Spearman’s *r*_*s*_=0.952; *R*^2^=0.879), as the larger sample results in a more highly resolved allele frequency spectrum. This increase in sample size does not increase either the time to train or to test NoDEAR relative to *n*=20 results.

### Robustness of recombination estimation from the allele frequency spectrum

The above tests were all carried out under relatively simple conditions. We used a machine learning model trained and tested on an equilibrium population history, using the derived allele frequency spectrum, at only one window size (50 kb), and with no gene conversion (which could bias estimates of *ρ*; Setter et al. 2022; Dutheil 2024). We therefore wanted to understand how robust our inferences were to additional model complexities.

We first trained and tested NoDEAR on the minor allele frequency spectrum, keeping all other conditions the same. As expected, this approach did worse than when using the derived spectrum, but only slightly so (Spearman’s *r*_*s*_=0.888; *R*^2^=0.701; Table S1). We also carried out training and testing using additional genomic locus sizes, including 10 kb, 100 kb, 150 kb, and 200 kb windows (keeping the per-base recombination rate the same in each). Across these window sizes, we found a range of correlations between the estimated and true values of *ρ* (Spearman’s *r*_*s*_: 0.847, 0.932, 0.940, and 0.944, respectively; Table S1). Although the smallest window size (10 kb) shows some reduction in predictive accuracy—possibly because there are not enough recombination events to distinguish among different values of *ρ*, or not enough SNPs to construct an accurate spectrum—NoDEAR behaves well at larger window sizes. However, we expect that at some larger window size predictive accuracy must go down, as every window will have an indistinguishably large number of recombination events and therefore indistinguishable values of *ρ*.

We assessed the effect of non-equilibrium demographic conditions by generating test data with such histories, but using a NoDEAR model that was trained on equilibrium conditions. We simulated two types of non-equilibrium demographies: in the first set, we sampled changes in population size through time by first drawing a number of times at which a population changed in size from a Poisson distribution with rate parameter, λ=3. This means that most simulated loci will have changed in size three times, but some will have changed two or four (or one or five, etc.) times as well. For each change in size, we then drew a new size from a normal distribution centered on *N*=70,000. These simulations therefore capture expansions and contractions in a single population over time, independently carried out for each locus. In the second set of non-equilibrium histories, we simulated population structure by having two sub-populations that split 10,000 generations ago for all loci (all populations have *N*=70,000). The allele frequency spectrum was constructed by sampling *n*=10 individuals from each of the two sub-populations and combining them for a total of *n*=20. In both sets of non-equilibrium simulations, the true value of *ρ* was again calculated via the number of recombination events in the history of a locus by using the relationship *ρ* =*R/a*.

Perhaps surprisingly (see Discussion), we observed no reduction in accuracy of NoDEAR when predicting *ρ* in populations whose sizes are changing over time, even when our model is trained on equilibrium histories. For 50-kb windows, results from test datasets of non-equilibrium histories are even a slightly bit better than with equilibrium histories (Spearman’s *r*_*s*_=0.932; *R*^2^=0.841; Figure 3b). Table S1 shows that while inference on non-equilibrium populations is not better across all window sizes, neither does it get much worse than results from equilibrium populations. Likewise, simulations with a history of population subdivision led to slightly worse results in 50-kb regions (Spearman’s *r*_*s*_=0.870; *R*^2^=0.778; Figure 3c), but no consistent reduction across other window sizes (Table S1).

Finally, we generated a test dataset in which both crossing-over and gene conversion can occur (all previous simulations only had crossing-over). Many LD-based methods for estimating *ρ* cannot capture the effects of both types of recombination, sometimes overestimating and sometimes underestimating (e.g. pyrho) the total recombination rate (Dutheil 2024). In these simulations gene conversion events make up 50% of all recombination events, with tract length 300 bp. Again, however, we see no reduction in the accuracy of NoDEAR, even though it was trained on data with no gene conversion (50-kb: Spearman’s *r*_*s*_=0.928; *R*^2^=0.738; Figure 3d; Table S1 contains all other window sizes). NoDEAR is accurately predicting the amount of recombination, regardless of the exact mechanism of recombination.

### Application to data from humans

To further demonstrate the use of the allele frequency spectrum as a method for estimating the population recombination parameter, we applied NoDEAR to a dataset from humans. Because we do not know the true recombination rates across loci in these data, we also ran pyrho for comparison. We chose 10 diploid samples from Finland (*n*=20), using only SNPs from chromosome 6, giving a total of 3,408 50-kb windows (The 1000 Genomes Project Consortium 2015).

One issue that occurs in real data that did not occur in our simulations is missing genotypes. To deal with missing data—which can result in different counts of the minor or derived allele at different sites—we used a vector with counts of SNPs in 10 bins of allele frequencies (e.g. 0-0.05, 0.05-0.10, etc.). Because alleles were not assigned as ancestral or derived in the human data, we used NoDEAR trained on the minor allele frequency spectrum. We only used 50-kb windows of the genome that contained at least 100 SNPs, to ensure that there was enough data for estimation. In total, this resulted in 2,602 50-kb windows that could be analyzed by NoDEAR.

NoDEAR ran quickly on all 2,602 loci, taking 25.4 seconds; in comparison pyrho took 55,100.2 seconds (i.e. 15.3 hours), including steps that inferred the population demography of the Finnish sample (which only took 156.4 seconds). The value of *ρ* inferred by NoDEAR varied from 0.237 to 367.6 across chromosome 6; pyrho reports *c* for the same data, ranging from 1.02×10^−10^ to 1.51×10^−7^ (taking the average of each window; we again removed 104 values with *c*≈10^−50^). Overall, there were 2,498 50-kb windows with estimates from both NoDEAR and pyrho. The correlation between the two estimates of recombination among these windows was quite high (Spearman’s *r*_*s*_=0.61; Supplementary Figure 3), providing further evidence that one can infer the recombination history of natural populations using only the allele frequency spectrum.

## Discussion

Recombination is of interest to a wide variety of biologists, but has an especially important role in evolutionary biology because of its role in moderating the influence of natural selection (e.g. Hill and Robertson 1966). As a result, there are many approaches for estimating recombination across the genome. One very common approach uses polymorphism data to estimate the amount of recombination in the history of a small sample, resulting in an estimate of the population recombination parameter, *ρ* (sometimes called *C*). This estimate represents an integrated history of recombination over time and across individuals that have left descendants in a sample.

Because most modern methods that estimate *ρ* use LD among sites, often population-based estimates of recombination are simply called “LD-based” estimates. It should be noted, however, that some of the first statistical estimators of *ρ* used the variance in pairwise differences between phased haplotypes, not LD (Hudson 1987; Wakeley 1997).

Regardless of the exact approach used, all previous model-based methods for estimating *ρ* required that the genotypes of the individuals being considered could be associated with those individuals. Sometimes individual alleles are arranged along chromosomes within individuals (i.e. gametic LD), and sometimes diploid genotypes along chromosomes are associated with individuals (i.e. genotypic or zygotic LD). Here, we have demonstrated that the allele frequency spectrum alone can be used to estimate population recombination rates. The allele frequency spectrum is a vector of SNP counts or proportions at each frequency in a sample, and contains no information about alleles or genotypes at different loci found in any particular individual.

However, we find that the allele frequency spectrum does indirectly contain information about the number of marginal gene trees in a region (Figures 1, 2). Because the number of gene trees reflects the number of recombination events, the spectrum can be used in a straightforward way to estimate *ρ*.

Our exploration of the role of the allele frequency spectrum in recombination estimation was inspired by the software ReLERNN (Adrion et al. 2020). ReLERNN is a machine learning method that can infer *ρ* from either genotype data or pooled sequencing data, carrying out both tasks with high accuracy. As pooled sequencing data does not contain any information on genotypes of individuals, we were curious about the “black box” at the heart of the ReLERNN machine learning model. We reasoned that the model was likely using the allele frequency spectrum to predict *ρ*, and indeed our analyses suggest that this is the case. Importantly, however, ReLERNN could be using additional information not considered by the simplified model learned by our software, NoDEAR. We purposefully removed information about both the number of SNPs in a window and the location of SNPs in each window. In the Introduction we discussed how the number of SNPs could be informative about *ρ* (at least in non-neutral scenarios), but the location of SNPs may be even more informative, including in neutral scenarios. One could imagine calculating the correlation of the allele frequency spectrum (or other measures of variation) among sub-windows of a larger region in order to see how quickly it changes. Such information is surely informative about *ρ*, and may be being used by ReLERNN. As our goal was largely to understand the information contained within the allele frequency spectrum—and neither to fully dissect ReLERNN nor to build a competitor software to it—we did not explore the possibilities contained within these other pieces of data further.

In addition to demonstrating that the allele frequency spectrum can be used to estimate recombination, we also showed that a very reduced representation of this spectrum could be used for the same purpose. We calculated the Euclidean distance (*L*^2^-norm) between the allele frequency spectrum from a region and the expected equilibrium spectrum, showing that this was highly predictive of *ρ* (Supplementary Figure 1). This simple summary statistic works well because it captures the main effect of recombination at a locus: that the spectrum generated by summing over many trees will be much smoother, and therefore closer in distance to the expected spectrum, than the spectrum from a small number of trees. We could of course have trained a machine learning model using this distance, but as it is a single value we would not do much better than the rank correlation between *L*^2^-norm and *ρ*, which was already quite high (Spearman’s *r*_*s*_=-0.89). How could one use *L*^2^-norm if a population did not have an equilibrium history? One straightforward solution is to build a reference allele frequency spectrum by summing over data from all loci together. This spectrum has the maximal amount of recombination possible, and therefore the Euclidean distance between individual windows and this reference should again be proportional to the recombination rate in each window.

One possibly surprising result from our simulations is that the inferences made by NoDEAR were not affected by non-equilibrium population histories—either changes in population size or structure—even when the model was trained on an equilibrium history. While non-equilibrium histories will have allele frequency spectra that have a different shape from the equilibrium spectrum produced during training, these results suggest that NoDEAR is not learning this shape per se, but rather the choppiness/smoothness of spectra generated by different levels of recombination. This behavior is quite helpful, as it means that one would not have to re-train the model on every new dataset.

Our results concerning the lack of an effect of non-equilibrium histories on the estimation of recombination stands in apparent contrast to some previous results, and require clarification. Multiple previous simulation studies have shown that non-equilibrium histories can affect estimates of the recombination rate (McVean et al. 2002; Smith and Fearnhead 2005; Johnston and Cutler 2012; Dapper and Payseur 2018; Spence and Song 2019; Adrion et al. 2020; Samuk and Noor 2022; Raynaud et al. 2023; Dutheil 2024). However, different studies often mean something slightly different by “recombination rate.” As mentioned earlier, *ρ* is the population recombination parameter, defined as 4*N*_e_*c*, such that both the population history at a locus and the per-generation recombination rate at a locus will affect the number of recombination events found in a sample. In an equilibrium population, estimates of *N*_e_ can be used to estimate *c* (sometimes called *r*) directly. Even in populations with non-equilibrium histories, accurate estimates of *N*_e_—taking into account this history—can be used to estimate *c* (Spence and Song 2019; Adrion et al. 2020). It is the estimate of *c* that can be biased when non-equilibrium histories are not taken into account (i.e. when the wrong value of *N*_e_ is used), not the amount of recombination that has occurred in a sample from nature (i.e. *ρ*).

Estimating *c* from *ρ* assumes that the inferred demographic model is the only cause of genealogical heterogeneity. If natural selection acts to either reduce (e.g. positive selection or background selection) or increase (e.g. balancing selection) the height of a genealogy, the number of recombination events may no longer be proportional to *c* (Smith and Fearnhead 2005; O’Reilly et al. 2008; Spence and Song 2019; Adrion et al. 2020). This occurs because the effective population size at such loci is not determined solely by demographic history. We have chosen to construct NoDEAR to only predict *ρ* from data; it is likewise trained only on *ρ*-values directly calculated from the actual number of recombination events produced by our simulations. It is this relatively assumption-free approach that allows NoDEAR to be accurate under different non-equilibrium conditions (and mechanisms of recombination, such as gene conversion) for which it was not trained. We imagine it would do a similarly accurate job of estimating *ρ* in the presence of non-neutral evolution.

The allele frequency spectrum is a fundamental measurement of variability in DNA sequences, used for the inference of both selection and demography. Here, we have shown that it also contains evidence of recombination, as it encodes information about the number of marginal gene trees in a genomic window. Although this property of the allele frequency spectrum has not been recognized before (that we are aware of), certainly there are many theoretical studies that are relevant to the utility of the spectrum for this purpose. For instance, work on the average distance between recombination events (e.g. Deng et al. 2021) and the average effect of recombination events on tree topologies (e.g. Ferretti et al. 2013) both provide important context for the power of the allele frequency spectrum alone to infer recombination. We hope that future work can further explore the application of this, or related, approaches.

## Data Availability Statement

All code used in this paper is available on GitHub (https://github.com/smishra677/NoDEAR).

## Acknowledgements

We thank Jeff Adrion and Andy Kern for discussion, the Kern-Ralph co-lab for helpful feedback at an early stage of this work, and Richard Wang for comments on the manuscript.

## Supplementary Figure legends

**Supplementary Figure 1.**
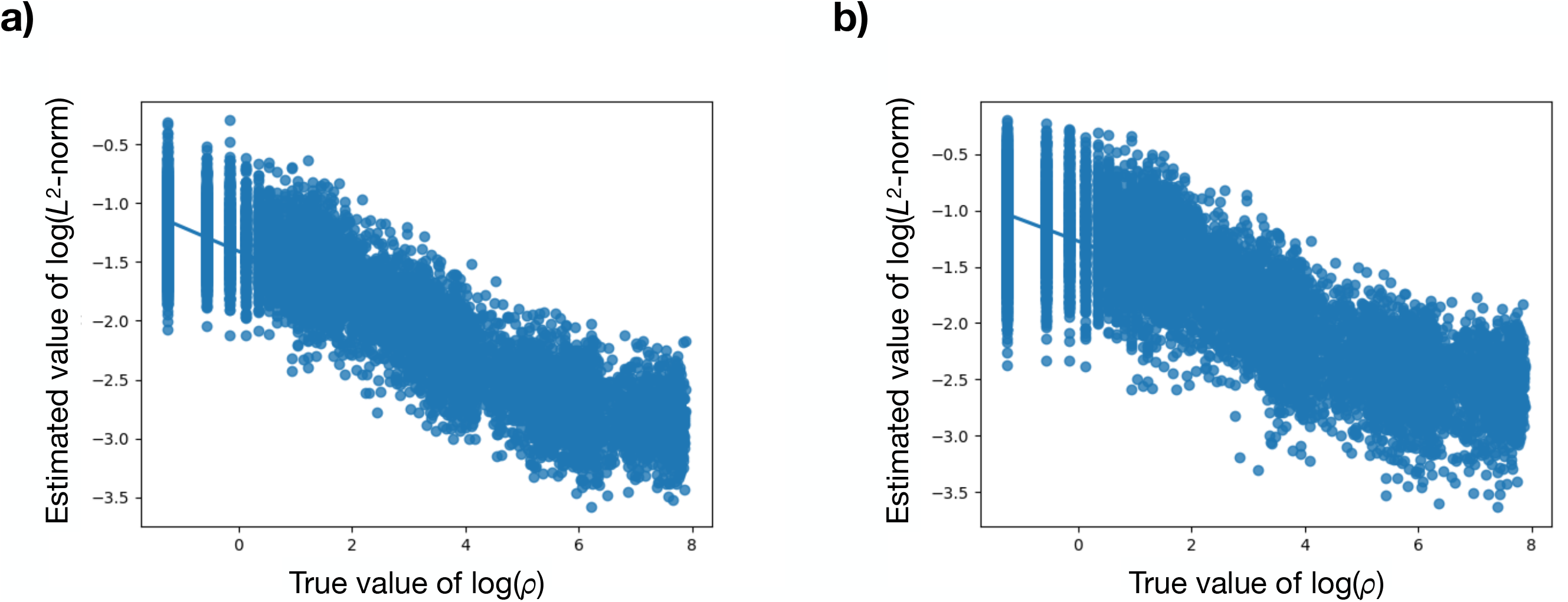
Correlation between true values of *ρ* and values of *L*^2^-norm calculated from allele frequency spectra. **a)** Correlation between *ρ* and values of *L*^2^-norm calculated from the derived allele frequency spectrum (Spearman’s *r*_*s*_=-0.89). Each dot represents one simulated 50-kb region. Blue lines represent best-fit regressions. **a)** Correlation between *ρ* and values of *L*^2^-norm calculated from the minor allele frequency spectrum (Spearman’s *r*_*s*_=-0.83).

**Supplementary Figure 2.**
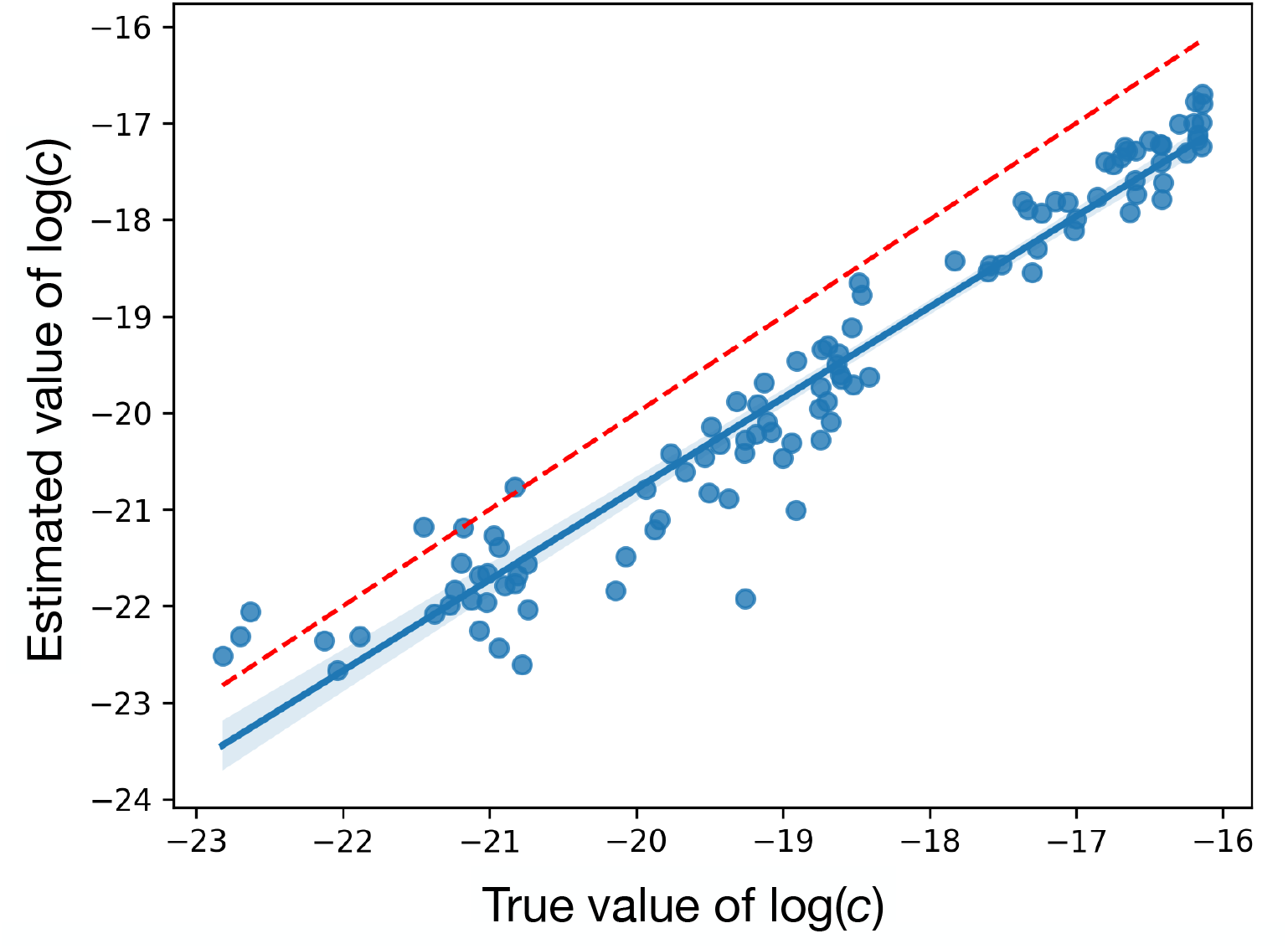
Correlation between true values of *c* and values estimated by pyrho (Spearman’s *r*_*s*_=0.97). Each dot represents one simulated 50-kb region The blue line represents the best-fit regression, plus confidence intervals. The red dashed line shows the *y*=*x* line.

**Supplementary Figure 3.**
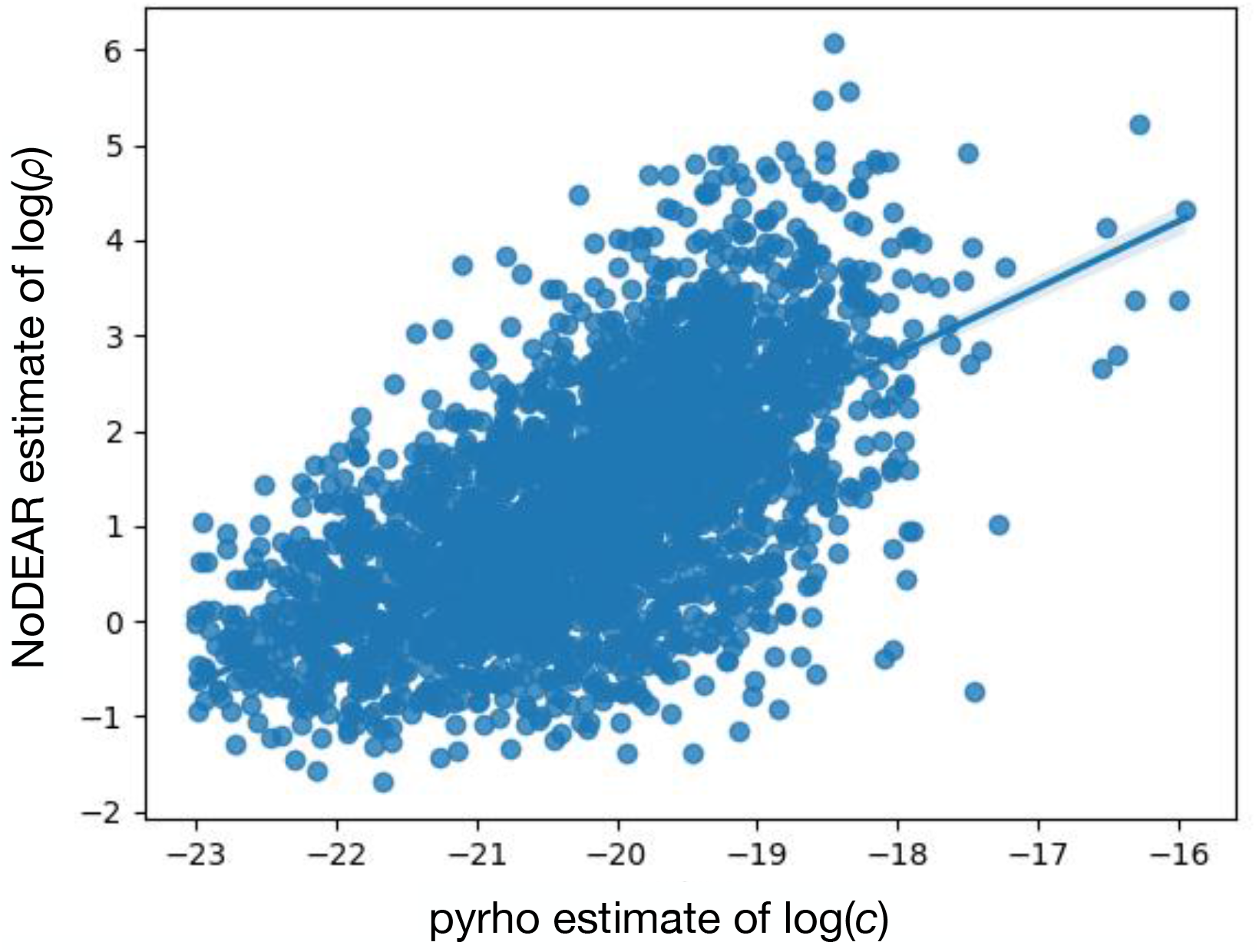
Correlation between values of *c* estimated by pyrho and values of *ρ* estimated by NoDEAR (Spearman’s *r*_*s*_=0.61). Each dot represents one 50-kb region of the human genome. The blue line represents the best-fit regression, plus confidence intervals.

